# Uropathogenic *Escherichia coli* population structure and antimicrobial susceptibility in Norfolk, UK

**DOI:** 10.1101/2023.03.24.533965

**Authors:** Cailean Carter, Alexandra Hutchison, Steven Rudder, Elizabeth Trotter, Emma Waters, Ngozi Elumogo, Gemma C. Langridge

## Abstract

**Background:** Half of all women have experienced a urinary tract infection (UTI) in their lifetime and this remains a persistent issue in rural counties like Norfolk, UK. In alignment globally, Uropathogenic *E. coli* (UPEC) are the main etiological agent for UTIs in Norfolk and are increasingly difficult to treat due to multi-drug resistance (MDR).

**Objective:** We set out to identify which clonal groups and resistance genes are disseminating in the community and hospitals in Norfolk, the first study of its kind for UPEC in this region.

**Methods:** We collected 217 clinical *E. coli* isolates causing UTIs in the community and hospital from the Clinical Microbiology laboratory at Norfolk and Norwich University Hospital. These were whole genome sequenced using the Illumina and MinION platforms for *in silico* multi-locus sequence typing and antibiotic resistance determinant detection.

**Results:** The isolates were composed of 74 sequence types (STs); 8 lineages represented 57% of this population: ST73, ST12, ST69, ST131, ST404, ST95, ST127, and ST1193. Importantly, primary UTI screening deemed 8% of isolates to be MDR, with high rates of resistance to ampicillin (52.1%) and trimethoprim (36.2%) in hospitals. Of concern is the probable clonal expansion of MDR groups ST131 and ST1193 in hospitals and community settings with chromosomally encoded *bla*_CTX-M-15_, *bla*_OXA-1_, and aac(6’)-Ib-cr5.

**Conclusions:** The burden of reported UTIs in Norfolk is largely caused by non-MDR isolates. The UPEC population is continually evolving, and monitoring samples with consideration of sources will help reduce burden of disease.

## Introduction

Urinary tract infections (UTIs) involving Uropathogenic *Escherichia coli* (UPEC) are a significant contributor for both visits to primary care providers and antibiotic prescribing in the UK. Besides monitoring antibiotic prescribing practices and antimicrobial resistance testing results for UTIs, England does not have comprehensive UTI surveillance to monitor national or regional UPEC populations.^1^ Therefore, independent regional studies report emerging multi-drug resistant (MDR) clonal groups using sequence typing (ST). One example is the emergence of ST131 in the early 2000s, arguably the most successful group in its frequency and acquisition of MDR determinants.^2^ The globally emerging ST1193 (first recorded in 2012) has gained interest internationally for acquisition of fluoroquinolone resistance from ST10 and subsequent clonal expansion.^3–5^

In this work, we investigated the UPEC population circulating in Norfolk, a rural county in East Anglia with a population of 916,200 people.^6^ In this region, 67.5% of reported UTIs are from women of all ages.^7^ However, the remaining third are reported by men, mostly between the ages of 66 and 85. This is significant as elderly men have been strongly associated with uncomplicated UTIs caused by MDR *E. coli* carrying extended-spectrum β-lactamases (ESBLs).^8, 9^ The expansion of MDR clonal groups limits therapeutic options and can mean deviating from cost-effective options. Such a trend may have contributed to the expected cost of £860,161 in 2021 for 88,459 antibiotic prescriptions for uncomplicated UTIs under NHS Norfolk and Waveney.^10^ Thus, we sought to better understand the landscape of UPECs being reported at community and hospital facilities by whole genome sequencing and antibiotic resistance phenotyping.

## Materials and methods

### Collection of *E. coli* isolates from UTI cases

A total of 217 clinical *E. coli* isolates from UTI patients were collected from the Norfolk and Norwich University Hospital (NNUH) Clinical Microbiology laboratory (Table S1). An initial 18 were collected in July 2020 and selected for their resistance profile to commonly prescribed antibiotics for UTIs: five trimethoprim resistant, five nitrofurantoin resistant, four trimethoprim and nitrofurantoin resistant, and four susceptible to both trimethoprim and nitrofurantoin. A further 199 were collected with source metadata between August 2021 and January 2022 irrespective of phenotype and patient attributes. Only samples that were positive for infection (> 35 white blood cells per μL and/or ≥ 9000 bacteria per μL) determined by the Sysmex UF 5000 Automated Microscopy system and 10^5^ colony forming units per litre on orientation (ORI) agar chromogenic plates were included. Repeat urine specimens were included during the collection period and records of repeat samples were kept. For each sample, a single purple colony on an ORI plate, confirmed to be *E. coli* and isolated from mid-stream and catheter specimens of urine was selected, streaked onto LB agar, and incubated overnight at 37°C. Subsequently, 1 mL of pre-warmed LB broth was added to each sample and incubated at 37°C overnight to prepare stocks in 20% glycerol that were stored at −80°C.

### Phenotypic resistance

Phenotypic resistance metadata was provided by NNUH. A first line urine profile was performed with ampicillin, cefalexin, cefpodoxime, ciprofloxacin, co-amoxiclav, gentamicin, nitrofurantoin, and trimethoprim using Oxoid discs following the latest EUCAST guidelines. Cefpodoxime was used for screening AmpC and/or ESBL production but is not part of the NHS formulary and is therefore not reported in this study.^11^ When the criteria for MDR was passed (resistance identified to nitrofurantoin and trimethoprim, ampicillin/co-amoxiclav and cefalexin, or cefpodoxime), isolates were put forward for further antibiotic screening on the Vitek 2 System (Biomérieux) with N351 Gram-negative susceptibility cards. Susceptibility to monobactams (piv)mecillinam and aztreonam was determined by disc diffusion for MDR isolates. Visualisation of phenotypic resistance was generated with Adobe Illustrator.

### Whole genome sequencing (WGS)

DNA extractions for the initial 18 isolates were performed with a GeneJET Genomic DNA Purification Kit (Thermo Scientific) following the manufacturer’s instructions. Fire Monkey DNA Extractions Kits (RevoluGen) were used for the remaining isolates, with the first 100 being performed manually using spin columns and remaining 109 being processed using Fire Monkey filters on a 96-well plate and positive air pressure on the Resolvex A200 robotic platform (Tecan). Manual DNA extracts were quantified using Qubit with a broad-range assay kit (Thermo Scientific); automated DNA extracts were quantified using Quant-IT broad range (Thermo Scientific) with a plate reader. Illumina sequencing was performed on the NextSeq as described by Parker and colleagues.^12^ Quality control of short reads were performed as described in Supp. Method 1.

### In-silico analysis

Most in-silico analyses were performed on the Quadram Institute cloud-based Galaxy server (release 19.05).^13^ Command-line tools were run on Ubuntu 20.04 LTS on Windows Subsystem for Linux 2. Python3 (v3.8.2) was used for data handling on Jupyter Notebooks with the pandas (v1.1.3) and NumPy (v1.18.3) packages. Fisher’s exact test and Chi-Squared Tests were performed using SciPy (v1.4.1). Unless otherwise stated, default parameters were used. Code used for this study can be found at https://github.com/CaileanCarter/UPEC_pop_study_Norfolk.

### Draft whole genome assemblies

Short read sequences were assembled with Shovill (v1.1.0)^14^ with down-sampling to average depth of 100x, estimated genome size of 5 Mbp, minimum contig length of 500 bp, minimum coverage of 2, and using the Spades assembler. Assembly completeness was inspected using BUSCO (v3.0.2)^15^ on the FASTA contig files.

### In silico typing

Sequence typing was performed using MLST (v2.16.1).^16, 17^ Phylogrouping was performed using EzClermont (v0.6.3).^18^ Plasmid typing by MLST was performed in-silico using plasmidfinder and matches were filtered for ≥95% identity and ≥95% coverage.^19^

### Core genome alignment, phylogeny, and clustering

To investigate the relationships in our dataset from the perspective of the core genome, a core genome alignment of short reads against *E. coli* UTI89 (NC_007946.1; GCF_000013265.1) was made using Snippy (Galaxy Version 4.4.3+galaxy2).^20^ On average, 84%±0.055 of the reference genome was aligned with sequencing reads. Snippy-core was used to combine the Snippy output into a single core SNP alignment for constructing a maximum likelihood (ML) phylogenomic tree with RAxML (v8.2.4).^21^ Visualisation and annotation was done in R (v4.2.1) using RStudio (v2202-07-22 for Windows). The R package APE (v5.6-2) was used to load the Newick file of the best scoring ML tree from RAxML.^22^ Plotting of the tree and metadata was done with ggtree (3.4.2) and ggplot2 (v3.3.6).^23–26^ Clustering of isolates were supported by the FastBAPS Bayesian hierarchical clustering algorithm using a core genome alignment and ML tree, and the ‘optimise.symmetric’ prior.^27^ Robustness of FastBAPS clustering was confirmed with a heatmap as described in the FastBAPS documentation. Any isolates not automatically assigned a phylogroup (n=4) or ST (n=15) were assigned to a plausible group by inference from the neighbouring isolates within the same subclade.

### Identification of AMR determinants and association with phenotypic resistance

Draft assemblies were annotated and translated into amino acid sequences using Prokka (v1.14.5).^28^ Annotations of AMR determinants was performed by AMRFinderPlus (v3.10.40) with inputs from Prokka and FASTAs of draft assemblies.^29^ As we could only identify a nitrofurantoin resistance determinant for 1 of 11 nitrofurantoin resistant isolates, a database of nitrofurantoin resistance determinants in *E. coli,* NITREc, was used.^30^ The UPEC amino acid sequences were aligned against this database using BLASTp (v2.10.1) and filtered for 100% identical matches and 100% positive-scoring matches. Association of resistance determinants with phenotypic resistance was calculated by Fisher’s Exact Test and corrected for false discovery rate using the Python statistics package statsmodels’ (v0.12.2) *fdrcorrection* method. To improve confidence of association, a Chi-Squared Test with Yates’ correction was used, and genes were filtered by statistical significance agreed by both tests. Strength of association, Cramér’s phi (*φ_c_*), was calculated from the Chi-Squared value (χ^2^) with the formula 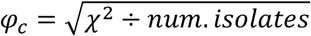. Relationship between source and antibiotic resistance phenotype was calculated by Chi-Squared Test with Yates’ correction. Isolates found to have the *bla*_OXA-1_, *bla*_CTX-M-1_, *bla*_CTX-M-15_, *bla*_SHV-1_, *bla*_DHA-1_, or aac(6’)-Ib-cr5 genes were long read sequenced via Oxford Nanopore Technologies (ONT) to confirm genomic location. Contigs where the *rep* gene was identified by plasmidfinder were cross-referenced with results from AMRFinderPlus for matching contig accessions, therefore connecting resistance genes with a given plasmid type.

### Native-ligation nanopore sequencing for AMR determinant location

Forty-eight samples were prepared in a single MinION library using the Ligation Sequencing Kit (SQK-LSK109, ONT) in conjunction with the Native Barcoding Expansion 96 (EXP-NBD196, ONT). The library was loaded onto the flow cell according to the manufacturer’s instructions. Sequencing was performed on the MinION platform using R9.4 flow cells (FLO-MIN106, ONT) with a run time of 48 hrs. ONT MinKNOW software (v4.0.5) was used to collect raw sequencing data and ONT Guppy (v6.3.8) was used for local base-calling, de-multiplexing and barcode trimming of the raw data.

### Genome assembly via Nanopore for AMR determinant location

Draft assemblies from Nanopore reads were made with the Flye assembler (Galaxy v2.9.0) with the following inputs: mode was set to ONT regular reads pre-Guppy5, 5 Mbp estimated genome size, 5 polishing iterations, and allowed to rescue short unassembled plasmids.^31^ A polishing step was performed on the draft assemblies with the long reads using Medaka (Galaxy v1.4.3) to generate a consensus sequence, r941_min_high_g303 was used as the model.^32^ Two round of polishing using long reads with Racon (Galaxy v1.3.1.1) was performed, minimap2 (Galaxy v2.17.2) was used to provide the alignment of long reads to draft assemblies for polishing.^33, 34^ Lastly, two rounds of polishing with short reads was performed with Pilon (Galaxy v1.20.1), also using minimap2 for alignment of short reads to draft assemblies.^35^ Each polishing stage was quality checked using CheckM (Galaxy v1.0.11) for assembly completeness, contamination, and strain heterogeneity.^36^

## Results

### Composition of *E. coli* causing UTIs in Norfolk

Using phylogroups as a broad typing method, *E. coli* from phylogroup B2 were responsible for most of the reported UTIs (70%) (Table 1). Other phylogroups each represented less than an eighth of the dataset with D (12.9%) and B1 (8.8%) being the second and third most common phylogroups, respectively. As isolates were found for all documented phylogroups of *E. coli*, this suggested that an evolutionarily diverse range of *E. coli* isolates were causing UTIs in Norfolk. Despite this diversity, a phylogenomic tree of the isolates collected revealed 8 clonal groups comprising 57.1% of the dataset with 7 of 8 in phylogroup B2 (Figure 1). We identified 74 STs in our dataset with ST73 as the most frequently isolated (Table 1). Of concern were MDR groups ST131 and ST1193 which were in the top 8 most frequently isolated clonal groups. Two isolates could not be assigned a ST by MLST. The frequencies of ST12, ST69, and ST131 in this study were very similar and likely equally contributed to presentations of UTIs during time of collection.

**Table 1.**
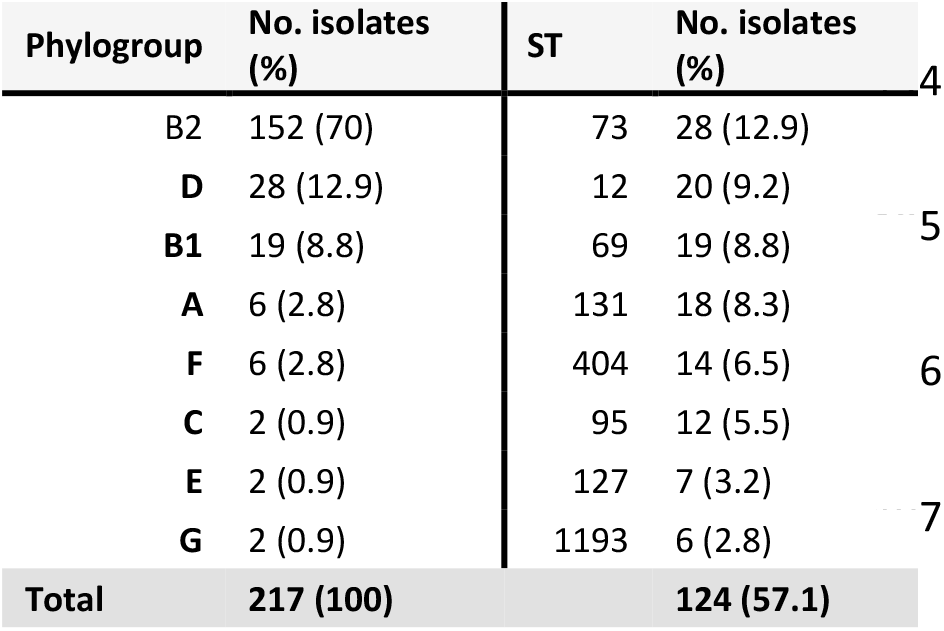
Phylogroups and predominant sequence types (STs)

**Figure 1.**
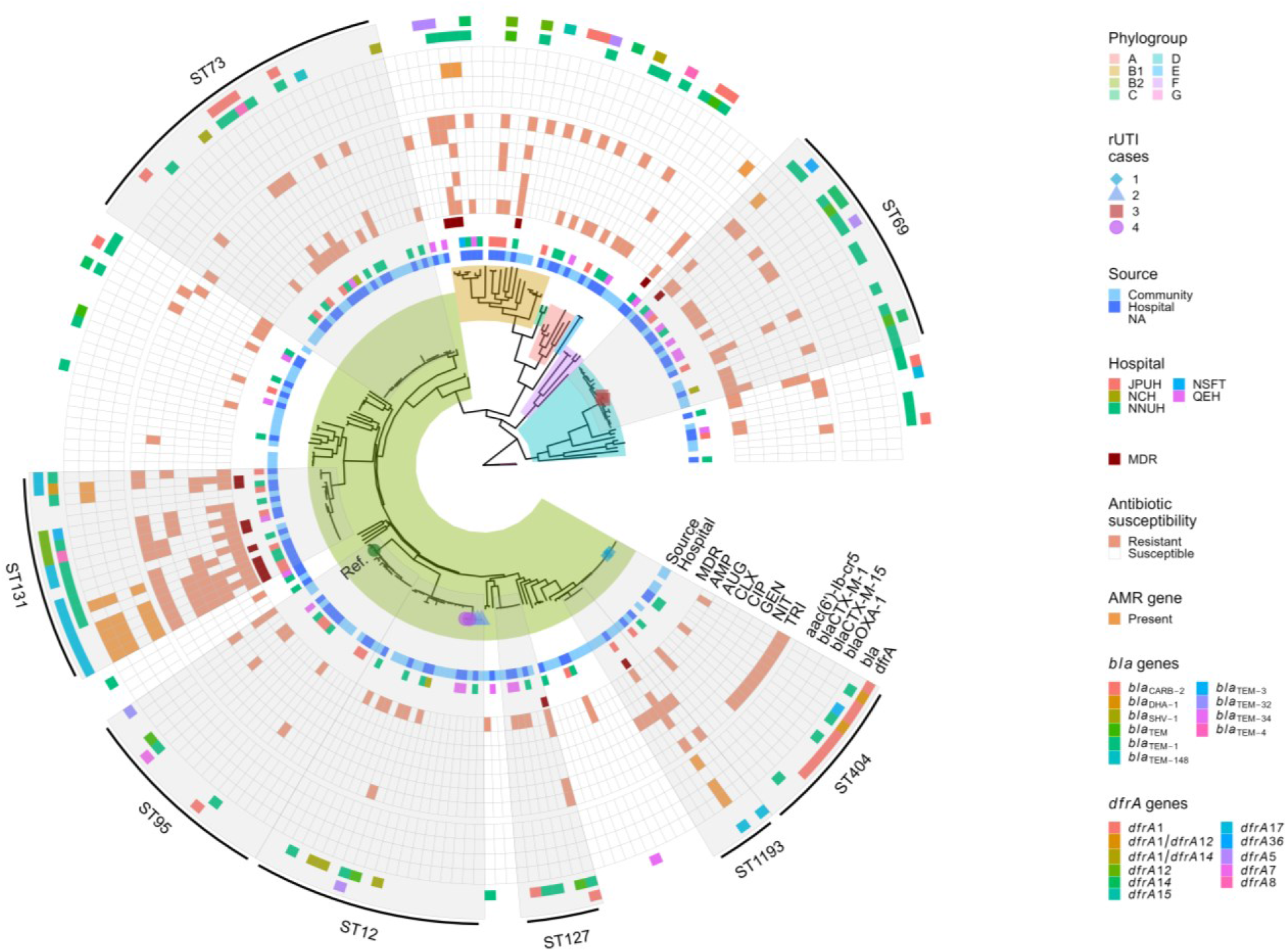
Phylogenomic tree of 217 *E. coli* isolates from UTI cases in Norfolk. Branch backgrounds are coloured by phylogroup. The reference genome, *E. coli UTI89*, is indicated by a dark green filled circle at the tip of the branch and labelled “Ref.” at the tip. Four rUTI cases are numbered and colour/shape coded at the branch tips. Metadata visualised include *source* (blank denotes isolates selected for their resistance profile towards nitrofurantoin and trimethoprim) and hospital of origin: James Paget University Hospital (JPUH), Norwich Community Hospital (NCH), NNUH, Norfolk & Suffolk Foundation Trust (NSFT) Waveny Ward, and Queen Elizabeth Hospital (QEH) King’s Lynn. Isolates classified as multi-drug resistant (MDR) and phenotypic antibiotic resistance (red) for ampicillin (AMP), co-amoxiclav (AUG), cefalexin (CLX), ciprofloxacin (CIP), gentamicin (GEN), nitrofurantoin (NIT), and trimethoprim (TRI). Followed by presence of aac(6’)-Ib-cr5, *bla*_CTX-M-1_, *bla*_CTX-M-15_, *bla*_OXA-1_, and other *bla* and all *dfrA* genes identified. Predominant STs are highlighted and labelled.

We collected isolates from hospital (n=94) and community (n=105) settings to determine and compare the *E. coli* composition reported in each setting. There was a roughly equal distribution of sources across the phylogenomic tree, including most predominant clonal groups. This suggests frequent movement between settings which is supported by the lack of clustering of isolates from a particular hospital (Figure 1). However, ST131 was largely sourced from hospitals (12/17 isolates; 73%) and ST1193 from the community (5/6 isolates; 83%).

Four patients provided multiple samples for this study. In each case, the sequencing of successive samples revealed patients were either reinfected with the same isolate or were unable to clear the infection between samplings. Interestingly, a branch deviated from the main ST12 cluster encapsulating 5 isolates from two separate recurrent (r)UTI cases (rUTI case #2 with 3 isolates, rUTI case #3 with 2 isolates) which has led to over-representation of ST12 in our dataset. Similarly, as 18 isolates were selected for their resistance profile, this could also create bias. Nonetheless, these isolates were distributed equally across the population and therefore did not significantly contribute to over-representation of a clonal group.

### Resistance rates for antibiotics used in UTI screening

For the purposes of analysing antibiotic resistance rates, the 18 isolates that were selected for their antibiotic resistance profile were excluded. A total of 199 isolates were used to explore antibiotic resistance trends. Overall, there was mostly broad susceptibility (>90% of isolates susceptible to a given antibiotic) to antibiotics used in UTI screening (Figure 2). The highest resistance rate in hospital and community settings was observed for ampicillin where just over half of hospital isolates (52.1%) and a quarter of community isolates (27.6%) were resistant to ampicillin. In contrast, the lowest resistance rate for hospital and community setting was observed for nitrofurantoin, the current first line treatment for uncomplicated UTIs. No clustering of nitrofurantoin resistance was observed but could be found in predominant clonal groups, suggesting relatively recent acquisition of nitrofurantoin resistance determinants (Figure 1). Despite a high resistance rate for trimethoprim in hospital (36.2%) and community (15.2%) settings, not all of the predominant ST groups harboured trimethoprim resistance (ST95 and ST127); in those that did the rate of resistance was highly variable (22-76%). Indeed, large clusters of trimethoprim resistance were observed in ST404 and ST131, with other groups exhibiting sporadic trimethoprim resistance gain/loss events (Figure 1). The overall resistance rate for ciprofloxacin was 9% and could be largely attributed to two groups: ST1193 and a subclade of ST131. Treatment options for ST131 were severely limited with two isolates resistant to all but one of the first line antibiotics used in UTI screening (with susceptibility to nitrofurantoin).

**Figure 2.**
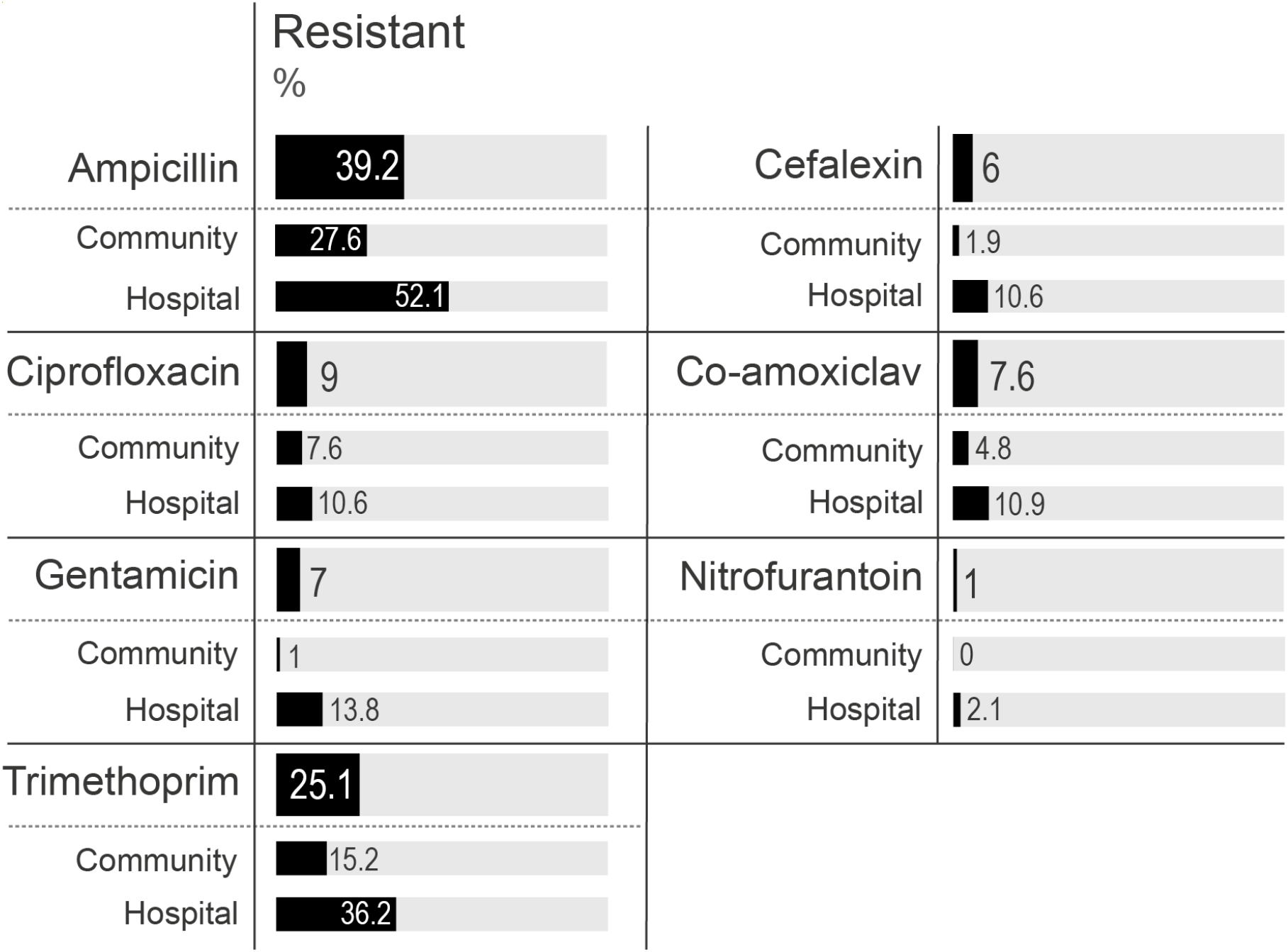
Antibiotic resistance rates. Percentage of isolates resistant to antibiotics used in first-line UTI treatment. Each antibiotic includes the total percentage on top and the percentage of isolates found in community and hospital resistant to that antibiotic below. (Total n=199, community n=105, hospital n=94)

We assessed the relationship between antibiotic resistance and source using a Chi-Square Test of Independence *X*^2^(1, N=199). This relationship was significant for ampicillin (*X*^2^ = 12.5, *p* = .0004), cefalexin (*X*^2^ = 6.67, p = .01), gentamicin (*X^2^* = 12.57, p = .0004), and trimethoprim (*X*^2^ = 11.5, p = .0007), evidencing higher resistance rates for these antibiotics in hospital settings. However, this relationship was not significant for co-amoxiclav (*X*^2^ = 2.46, p = .117) and ciprofloxacin (*X*^2^ = 0.55, p = .46). The test could not be performed on nitrofurantoin due to insufficient number of expectants.

### AMR determinants associated with phenotypic resistance

A curated database of known resistance mechanisms was used to identify resistance genes and point mutations associated with phenotypic antibiotic resistance within our dataset. Fisher’s exact test was used to determine the statistical significance of gene association with a given antibiotic resistance trait (p<0.05) and the strength of the association was determined with Cramér’s phi which provides a numerical scale of weak (0) to strong (1) association. To increase the scope of resistant determinants found in UTI cases, the 18 isolates selected for their antibiotic resistance profile were included in this analysis.

Of the total 18 antibiotics screened in this study (including the additional screening for MDR isolates), 14 were β-lactams – emphasising the importance of this family of antibiotics for UTI treatment. Thus, we assessed the association between (extended-spectrum) β-lactamase encoding genes with phenotypic resistance to β-lactams. Thirteen different β-lactamase determinants were detected across 87 β-lactamase-carrying isolates (Figure 1). Of these, *bla*_TEM-1_ was the most frequently identified β-lactamase (n=59) and was strongly associated with ampicillin resistance (p < .005) (Table 2). However, isolates found with just the *bla*_TEM-1_ β-lactamase had inconsistent resistance profiles against penicillins and co-amoxiclav, suggestive of one or more undetermined resistance mechanisms. The most frequently identified ESBL determinant was *bla*_CTX-M-15_ (n=11) which was strongly associated with cefalexin resistance (p < .005, 0.86 Cramér’s phi). Indeed, 8/11 isolates carrying *bla*_CTX-M-15_ were resistant to all 5 cephalosporins tested (Table S1). This gene was predominantly found in MDR groups ST131 (n=7) and in ST1193 (n=1), but also in ST85 (n=2) and ST3177 (n=1). Concerningly, *bla*_CTX-M-15_ frequently co-occurred with *bla*_OXA-1_, reducing susceptibility to β-lactam/inhibitor combinations, and aac(6’)-Ib-cr5, conferring resistance to aminoglycosides and fluoroquinolones.^37^ Through long read assembly, we verified the location of *bla*_CTX-M-15_, *bla*_OXA-1_, and aac(6’)-Ib-cr5 in ST131 (n=4) and ST1193 (n=1) to be chromosomally encoded (Table S3). A diversity of co-amoxiclav resistance determinants were identified but most were not statistically associated, this included *bla*_TEM-30_, *bla*_TEM-32_, *bla*_TEM-34_, *bla*_TEM-40_, and *bla*_TEM-148_ (Figure 1).^38^ Screening of carbapenem resistance was performed for a cohort of MDR isolates and carbapenem resistance was not detected in this study. However, one isolate from ST127 was found to carry a carbapenemase, *bla*_CARB-2_, which was not screened for carbapenem resistance as the isolate was deemed only ampicillin resistant in preliminary screening. Other β-lactamases detected in this study include *bla*_SHV-1_ (n=5), *bla*_CTX-M-1_ (n=1), and *bla*_DHA-1_ (n=1).

**Table 2.** Association of antibiotic resistance genes and point mutations with phenotypic resistance. Genes are listed with phenotypes with statistically significant association. Phenotypes are resistance to ampicillin (AMP), co-amoxiclav (AUG), ciprofloxacin (CIP), cefalexin (CLX), gentamicin (GEN) and trimethoprim (TRI). The combinations of genotype (G) and phenotype (P) presence/absence (+/-) are listed which were used for input for Fisher’s exact test (p-value corrected and odds ratio). Cramér’s association was used to determine strength of associations. Inf denotes an infinite value resulting from division of 0.

| Gene | Phenotype | G+ P+ | G- P+ | G+ P- | G- P- | p-value corrected | Cramér assoc. | Odds ratio |
| --- | --- | --- | --- | --- | --- | --- | --- | --- |
| *bla*<sub>TEM-1</sub> | AMP | 55 | 30 | 4 | 128 | 8.75E-23 | 0.68 | 59 |
| *bla*<sub>CTX-M-15</sub> | AMP | 10 | 75 | 1 | 131 | 1.89E-03 | 0.24 | 17 |
| *bla*<sub>OXA-1</sub> | AMP | 6 | 79 | 0 | 132 | 1.11E-02 | 0.21 | inf |
| *bla*<sub>SHV-1</sub> | AMP | 5 | 80 | 0 | 132 | 2.57E-02 | 0.19 | inf |
| *bla*<sub>TEM-40</sub> | AUG | 2 | 16 | 0 | 199 | 2.04E-02 | 0.32 | inf |
| *gyrA* D87N | CIP | 15 | 3 | 0 | 199 | 5.66E-19 | 0.91 | inf |
| *parC* S80I | CIP | 15 | 3 | 2 | 197 | 4.99E-17 | 0.85 | 493 |
| *gyrA* S83L | CIP | 16 | 2 | 13 | 186 | 2.01E-13 | 0.67 | 114 |
| *parC* E84V | CIP | 7 | 11 | 0 | 199 | 6.24E-08 | 0.61 | inf |
| *parE* L416F | CIP | 6 | 12 | 0 | 199 | 8.98E-07 | 0.56 | inf |
| *aac(6')-Ib-cr5* | CIP | 6 | 12 | 0 | 199 | 8.35E-07 | 0.56 | inf |
| *parE* I529L | CIP | 7 | 11 | 11 | 188 | 6.61E-04 | 0.33 | 11 |
| *bla*<sub>CTX-M-15</sub> | CLX | 10 | 2 | 1 | 204 | 2.01E-13 | 0.86 | 1020 |
| *bla*<sub>OXA-1</sub> | CLX | 6 | 6 | 0 | 205 | 6.15E-08 | 0.70 | inf |
| *aac(3)-IIe* | GEN | 6 | 9 | 0 | 202 | 2.66E-07 | 0.62 | inf |
| *aac(3)-IId* | GEN | 7 | 8 | 3 | 199 | 1.03E-06 | 0.55 | 58 |
| *aac(6')-Ib-cr5* | GEN | 5 | 10 | 1 | 201 | 2.50E-05 | 0.51 | 101 |
| *aadA5* | GEN | 6 | 9 | 5 | 197 | 7.29E-05 | 0.43 | 26 |
| *aadA2* | GEN | 4 | 11 | 5 | 197 | 5.87E-03 | 0.31 | 14 |
| *dfrA1* | TRI | 22 | 37 | 3 | 155 | 1.81E-10 | 0.49 | 31 |
| *dfrA17* | TRI | 14 | 45 | 0 | 158 | 3.56E-08 | 0.43 | inf |
| *dfrA12* | TRI | 7 | 52 | 0 | 158 | 4.02E-04 | 0.30 | inf |
| *dfrA5* | TRI | 5 | 54 | 0 | 158 | 5.23E-03 | 0.25 | inf |
| *dfrA14* | TRI | 6 | 53 | 1 | 157 | 6.59E-03 | 0.24 | 18 |

Ciprofloxacin resistance in our dataset was mostly attributed to ST131 and ST1193. This resistance was strongly associated with point mutations in *gyrA* D87N, *gyrA* S83L, and *parC* S80I which co-occurred in the same 15 ciprofloxacin resistant isolates (Table 2). In addition to point mutations in *gyrA* and *parC*, ciprofloxacin resistant ST131 isolates also carried the aac(6’)-Ib-cr5 gene which was significantly associated with both ciprofloxacin and gentamicin resistance. Trimethoprim resistance was associated with one of several *dfrA* derivatives with 6% of resistant isolates having 2 variations of *dfrA.* The most common was *dfrA1* which was found in 37% of trimethoprim resistant isolates. Nitrofurantoin resistance mechanisms were not identified using the above approach, therefore, a specialised database was used. Three of eleven nitrofurantoin isolates were found to have known resistance mechanisms through point mutations in *nfsA* and *nfsB* (exact point mutations were not specified by the database). Therefore, most nitrofurantoin resistance in our dataset could not be explained with existing databases.

## Discussion

The composition of predominant STs causing UTIs in Norfolk is reflective of predominant groups causing UTIs and bacteraemia in similar reports across the UK, Europe, and United States but in varying proportions.^2, 39–46^ Many of these STs are associated with animal sources (including food, wild, and companion animals) as detailed by Riley^47^ and Vincent *et al.,^48^* and can persist in the human gut.^49, 50^ Largescale studies of UK ESBL-producing *E. coli* imply ST131 is largely associated with humans, faeces, and sewage sources.^51, 52^ Indeed, some UK rivers have been identified as contaminated with CTX-M carrying ST131, thus, it is worth considering the contribution of Norfolk’s rivers and Broads as potential sources of ST131.^53^ Although there are reports of ST131 isolated from chickens, the subclade of ST131 (namely H22) attributed to this only accounted for 1 of 18 ST131 isolates collected in this study, suggesting poultry as an unlikely source of UTIs caused by ST131 in Norfolk.^51, 52, 54^ Given the recent emergence of ST1193, the potential sources of this group remain unclear.^3^ A recent phylogenomic analysis of 707 publicly available ST1193 sequences offer evidence to suggest companion dogs, urban-adapted birds, and wastewater as potential sources.^55^ Within ST131 and ST1193, all isolates carrying *bla*_CTX-M-15_, *bla*_OXA-1_, and aac(6’)-Ib-cr5 that were long read sequenced were found to reside on the chromosome; this has been reported globally with some suggesting this is indicative of another clonal expansion of ST131.^56–59^ We are in agreement with Ludden *et al.,^59^* that given the endogenous source of ST131 (and plausibly ST1193), measures should be taken to limit endogenous infections. Interestingly, the broadly susceptible ST12 group has not previously appeared as a predominant causative agent of UTIs in the UK, however, has been observed in Spain, France, and US and causing bacteraemia in the UK.^41, 46, 60, 61^ A limitation of this study is that we did not sample from surrounding environments, animals, or foods to identify sources of UPEC isolates in Norfolk. Nonetheless, these results suggest Norfolk is no exception to the global movement of UPEC given its similarity to the rest of the UK and international populations.

Since 2014, trimethoprim resistance in UPEC in Norfolk hospitals marginally reduced from 40% to 36.2%, whereas community rates halved from 33% to 15.2% possibly reflecting the change in antibiotic prescribing practices to nitrofurantoin as a first-line drug.^7^ Our reported resistance rates for trimethoprim closely match the reported rates by NHS Norfolk & Waveney CCG for 2021 Q3 to 2022 Q1.^62^ The overall resistance rate for nitrofurantoin is a third of the rate observed across the UK during the period of collection.^62^ Thus, current nitrofurantoin and trimethoprim resistance trends reported here and by the UK Health Security Agency suggest suitable efficacy in this region as first-line treatment for UTIs using current regional NICE guidelines. We were only able to identify a means of nitrofurantoin resistance in 3 of 11 nitrofurantoin resistant isolates via point mutations in *nfsA* and *nfsB*. Given nitrofurantoin is the current empirical treatment for UTIs, effective means of monitoring resistance determinants for nitrofurantoin are still needed as resistance rates are expected to increase. Ciprofloxacin has been used frequently as prophylaxis for prostate biopsies in Norfolk.^63^ However, the prevalence of ciprofloxacin resistance attributed to global MDR groups ST131 and ST1193, as seen here, raises concerns about the antibiotic’s efficacy as prophylaxis.^64^

In conclusion, the population of UPEC causing UTIs in Norfolk mirrors UPEC populations reported nationally and internationally. The ongoing evolution and presence of predominant clonal groups limits cost-effective treatment options for many UTI cases and continued monitoring can inform policy making to limit the burden of disease.

## Acknowledgements

We would like to thank Leonardo de Oliveira Martins, John Wain and Lisa Crossman from the Quadram Institute for their technical support and recommendations. Further, we thank Rhiannon Evans and David Baker of the Quadram Institute for their assistance with whole-genome sequencing. We would also like to thank the Clinical Microbiology lab at NNUH, especially Stephanie Walker, for their time and assistance in collecting isolates. Lastly, thank you to RevoluGen for providing FireMonkey HMW DNA Extraction Kits free of charge for this study.

## Transparency declaration

SR PhD study is partially supported by RevoluGen who provided FireMonkey HMW DNA Extraction Kits free of charge for this study but had no further input. GCL has previously consulted for RevoluGen on bioinformatic analyses. Authors declare no further conflicts of interest.

## Supplementary

### Supp. Method 1: Quality control of short-read data

All short reads were quality controlled using fastp (v0.19.5) and visualised using MultiQC (v1.11) to check reads were adequate for assembly and genotyping.^65, 66^ Read sets were deemed good quality with <5% duplication rate before filtering (low number of duplicate reads), GC content 50±5%, and >90% reads passing the quality and length filter. To assert an adequate number of reads were available for draft genome assembly, isolates were checked for short read coverage against the reference genome of *Escherichia coli* UTI89 (NC_007946.1; GCF_000013265.1) using samtools depth for BAM files with Snippy. If coverage was <30X, those isolates were re-sequenced and the reads merged with the previous run. QC summary can be found in Table S2. Isolates were again confirmed as *E. coli* using RefSeq Masher Matches (v0.1.1) by asserting the top match was *E. coli.^67^*

**Table S2.** QC summary statistics for short reads (n=217)

| Stat | Mean | Q75 | Q50 | Q25 |
| --- | --- | --- | --- | --- |
| **N50** | 213 Kbp | 267 Kbp | 209 Kbp | 153 Kbp |
| **No. contigs** | 144 | 111 | 86 | 69 |
| **Largest contig** | 0.5 Mbp | 0.64 Mbp | 0.52 Mbp | 0.39 Mbp |

**Table S3.** Summary of genomic location of antibiotic resistance genes of interest as determined by short read (SR) and located using long read (LR) sequencing. Plasmid type is the closest matching type by % identity and % coverage against reference *rep* genes. NI: not identified.

| ID | ST | Genes found by SR | Location by LR |  | No. contigs | Strain heterogeneity (%) | Contamination (%) | Plasmid type (% iden., % cov.) |
| --- | --- | --- | --- | --- | --- | --- | --- | --- |
|  |  |  | Chromosome | Plasmid |  |  |  |  |
| GL127 | 131 | aac(3)-Ile<br>aac(6')-Ib-cr5<br>*bla*<sub>OXA-1</sub><br>*bla*<sub>CTX-M-15</sub><br>*aadA5*<br>*dfrA17* | aac(3)-Ile<br>aac(6')-Ib-cr5<br>*bla*<sub>OXA-1</sub><br>*bla*<sub>CTX-M-15</sub> | *aadA5*<br>*dfrA17* | 5 | 0 | 0.33 | IncFIA (99.5, 99.7) |
| GL146 | 3177 | *bla*<sub>CTX-M-15</sub> |  | *bla*<sub>CTX-M-15</sub> | 8 | 9.09 | 0.45 | IncFIA (99.7, 100) |
| GL153 | 131 | *bla*<sub>CTX-M-15</sub><br>*bla*<sub>TEM-1</sub> |  | 1: *bla*<sub>CTX-M-15</sub><br>2: *bla*<sub>TEM-1</sub> | 3 | 20 | 0.89 | 1: IncFIB (99.7, 100)<br>2: IncFII (99.6, 100) |
| GL163 | 131 | aac(3)-Ile<br>aac(6')-Ib-cr5<br>*bla*<sub>OXA-1</sub><br>*bla*<sub>CTX-M-15</sub><br>*aadA5*<br>*dfrA17* | aac(3)-Ile<br>aac(6')-Ib-cr5<br>*bla*<sub>OXA-1</sub><br>*bla*<sub>CTX-M-15</sub> | *aadA5*<br>*dfrA17* | 12 | 0 | 0.33 | IncFIA (99.5, 99.7) |
| GL164 | 1193 | aac(3)-Ile<br>aac(6')-Ib-cr5<br>*bla*<sub>OXA-1</sub><br>*bla*<sub>CTX-M-15</sub><br>aph(3'')-Ib<br>aph(6)-Id<br>*dfrA17* | aac(3)-Ile<br>aac(6')-Ib-cr5<br>*bla*<sub>OXA-1</sub><br>*bla*<sub>CTX-M-15</sub> | *aph(3'')-Ib*<br>*aph(6)-Id*<br>*dfrA17* | 2 | 0 | 0.39 | IncFIA (99.5, 99.7) |
| GL172 | 69 | *bla*<sub>TEM-1</sub><br>*bla*<sub>CTX-M-1</sub><br>aph(3'')-Ib<br>aph(6)-Id<br>*aadA2*<br>*dfrA5* |  | *bla*<sub>TEM-1</sub><br>*bla*<sub>CTX-M-1</sub><br>aph(3'')-Ib<br>aph(6)-Id<br>*aadA2*<br>*dfrA5* | 2 | 0 | 0.09 | IncFII (100, 100) |
| GL183 | 131 | *aac*(3)-Ile<br>*aac*(6')-Ib-cr5<br>*bla*<sub>OXA-1</sub><br>*bla*<sub>CTX-M-15</sub><br>*aadA5*<br>*dfrA17* | *aac*(3)-Ile<br>*aac*(6')-Ib-cr5<br>*bla*<sub>OXA-1</sub><br>*bla*<sub>CTX-M-15</sub> | *aadA5*<br>*dfrA17* | 3 | 0 | 0.33 | IncFIA (99.5, 99.7) |
| GL190 | 12 | *aadA1*<br>*bla*<sub>SHV-1</sub> | *bla*<sub>SHV-1</sub> | *aadA1* | 3 | 0 | 0.12 | IncFIB (99.6, 100) |
| GL204 | 131 | *aac*(3)-Ile<br>*aac*(6')-Ib-cr5<br>*bla*<sub>OXA-1</sub><br>*bla*<sub>CTX-M-15</sub><br>*aadA5*<br>*dfrA17* | *aac*(3)-Ile<br>*aac*(6')-Ib-cr5<br>*bla*<sub>OXA-1</sub><br>*bla*<sub>CTX-M-15</sub> | *aadA5*<br>*dfrA17* | 2 | 0 | 0.33 | IncFIA (99.5, 99.7) |
| GL239 | 58 | *bla*<sub>CTX-M-15</sub> | NI | NI | 2 | 0 | 0.57 |  |
| GL240 | 58 | *bla*<sub>CTX-M-15</sub> | NI | NI | 9 | 0 | 0.23 |  |

